# Cortical stroke produces secondary injury and long-lasting gliosis in the ipsilateral thalamus

**DOI:** 10.1101/2020.12.24.424329

**Authors:** Gab Seok Kim, Jessica M. Stephenson, Abdullah Al Mamun, Ting Wu, Monica G. Goss, Jia-Wei Min, Jun Li, Fudong Liu, Sean P. Marrelli

## Abstract

Remote secondary injury in the thalamus has been observed following cortical infarct, however the mechanisms are not well understood. We used the distal MCAO stroke model (pdMCAO) to explore the cellular and temporal gliosis response in secondary thalamic injury in mice. At 3 days post-stroke (PSD3), primary infarct was limited to the cortex, with no infarct in the thalamus. However, at 2 weeks after stroke (PSD14), the ipsilateral thalamus demonstrated degenerating and severely damaged neurons. Staining for GFAP (astrogliosis) or IBA-1 (microgliosis) was first apparent in the ipsilateral thalamus by PSD3, and showed a progressive increase through PSD14. The number of activated microglia was increased within the thalamus at PSD14, reflecting proliferation of resident microglia as well as infiltration of peripheral monocytes. Interestingly, astrogliosis within the thalamus was enduring, as it was still evident at two years post-stroke. Furthermore, the astrogliosis at two years (but not at 6 weeks) demonstrated glial scar-like characteristics. Lastly, we demonstrated that post-stroke treatment with an NMDA receptor antagonist (memantine) reduces gliosis in the thalamus at PSD14. These findings highlight the development of lasting secondary injury in the thalamus following cortical stroke and support the value of memantine treatment in the mitigation of this injury.

## Introduction

It is a well-accepted concept that ischemic damage can spread from the initial damage site (primary injury site) to remote areas which are functionally linked with the primary injury region. These remote lesions can develop over time, which trigger a variety of cellular responses in specific brain regions ^1, 2^. Various cellular and molecular responses and injuries that aggravate neuronal and non-neuronal cell death are present in certain regions after stroke. These responses and changes in metabolic demand, neuro-inflammation, and cell death are mediated by excitotoxicity, reactive oxygen species (ROS) generation, apoptosis, necrosis, inflammations, glial cell activation, lymphocyte infiltration and blood brain barrier disruption, finally leading to brain infarction ^1, 3, 4^.

The thalamus has received little attention in the fields of stroke and neurodegenerative diseases, despite the thalamus’ critical role in relaying signaling between different brain regions (e.g. cortex, hippocampus) as well as the spinal cord ^5^. In addition, the thalamus is a key component in Papez circuits that are responsible for memory and cognition ^5, 6^. In fact, remote secondary thalamic injury has been observed in human stroke ^7, 8^ and has been detected in rodent stroke model ^9, 10^. Multiple pieces of evidence suggest that this thalamic injury contributes to worsened outcome and recovery ^11, 12^. Anatomically, the somatosensory cortex is connected to the thalamus via cortico-thalamic and thalamo-cortical projections ^13–15^. The cortico-thalamic projections which terminate in the somatosensory cortex originate from the ventral posterior nucleus of the thalamus and posteromedial complex (PoM)^16, 17^. This region is generally regarded as a control center for bidirectionally relaying signals between cerebral somatosensory cortex and peripheral nerves and is critical for several brain functions, such as consciousness, sleep, and respiration ^18–20^. Moreover, emerging lines of evidences clearly show that the thalamus is not a mere passive relay center, but also serves as a critical hub for integrating and relaying the multimodal signals from other brain areas ^21^. Dysfunction in the thalamus, caused by primary damage in cortex, is closely associated with poor outcome, degree of the disability and even decline of brain functions, such as cognition and memory ^22^. For these reasons, secondary thalamic injury would reasonably be expected to impair recovery following stroke.

Recently, our studies as well as others have shown that neuronal damage can be observed in thalamus, along with severe gliosis and increased inflammation in an experimental stroke model ^23–25^. It has been speculated that many factors including ROS, excitotoxicity, and inflammation that is amplified by glial cell activation in a damaged cortex may provoke and initiate secondary injury in the thalamus ^10^. However, detailed mechanisms and molecular changes in the thalamic area whereby cortical infarction leads to delayed injury in non-adjacent distant regions of brain are currently not well understood. We further sought to test the applicability of NMDA receptor antagonist (memantine) as a means for modulating secondary injury and gliosis in thalamus after cortical stroke. Also, as stroke predominantly occurs in the aged population, we additionally evaluated secondary injury in aged mice.

Here, we present data, which shows time-dependent molecular and cellular changes in the thalamus after stroke, focusing on neuronal loss and gliosis using a well-established stroke model in which primary infarct is limited to the cortex (i.e. no infarct in thalamus). We demonstrated monocyte infiltration as part of the thalamic pathology as well as microglial activation following stroke. We show that astrogliosis develops several days after primary cortical injury and can transition over time into a glial scar, which may persist permanently. In addition, memantine, NMDA antagonist, mitigated these responses in the thalamus. Together, our data suggests that reactive astrocytes and activated microglia are a significant component of secondary injury in the thalamus. To our knowledge, this study represents the first demonstration that 1) primary cortical injury induces microglial proliferation and monocyte infiltration in the thalamus, and 2) that astrogliosis in the thalamus is likely permanent. Lastly, we demonstrate that memantine treatment in the post-stroke period attenuates secondary thalamic injury. These studies thus provide further molecular evidence supporting secondary injury as an important component of stroke pathology and provide new mechanistic understanding from which to design future therapeutic strategies.

## RESULTS

### Cortical stroke induces neurodegeneration in ipsilateral thalamus

The <u>p</u>ermanent <u>d</u>istal <u>M</u>iddle <u>C</u>erebral <u>A</u>rtery <u>O</u>cclusion (pdMCAO) model was employed as an experimental model to explore the temporal development of gliosis associated with secondary injury in the thalamus. We first verified that the primary infarct was indeed restricted to the cortex, and specifically *not* evident in other brain regions, at a time point where primary injury is mature (PSD 3). Fig 1A shows representative TTC stained sections from a young C57BLJ mouse demonstrating well developed infarct in the cortex (white region), but not in the ipsilateral thalamus, hippocampus, or other brain regions within the striatum. We next used Fluoro-Jade C (FJC) staining to look for delayed neuronal degeneration within the thalamus after stroke. At PSD 14, FJC staining showed clear evidence of delayed neuronal injury in the ipsilateral thalamus, whereas the contralateral thalamus showed no increased staining (Fig. 1B). Additional analysis by cresyl violet (CV) staining further showed the presence of degenerating neurons, as evidenced by cell body shrinkage and darkly stained pyknotic nuclei, which are hallmarks for neuronal damage via pyknosis (Fig 1C,D). Taken together, these data indicate 1) that cortical infarction leads to neuronal damage in the thalamus in the late stroke phase and 2) that the pdMCAO stroke model provides a good model for evaluating secondary injury following stroke.

**Figure 1.**
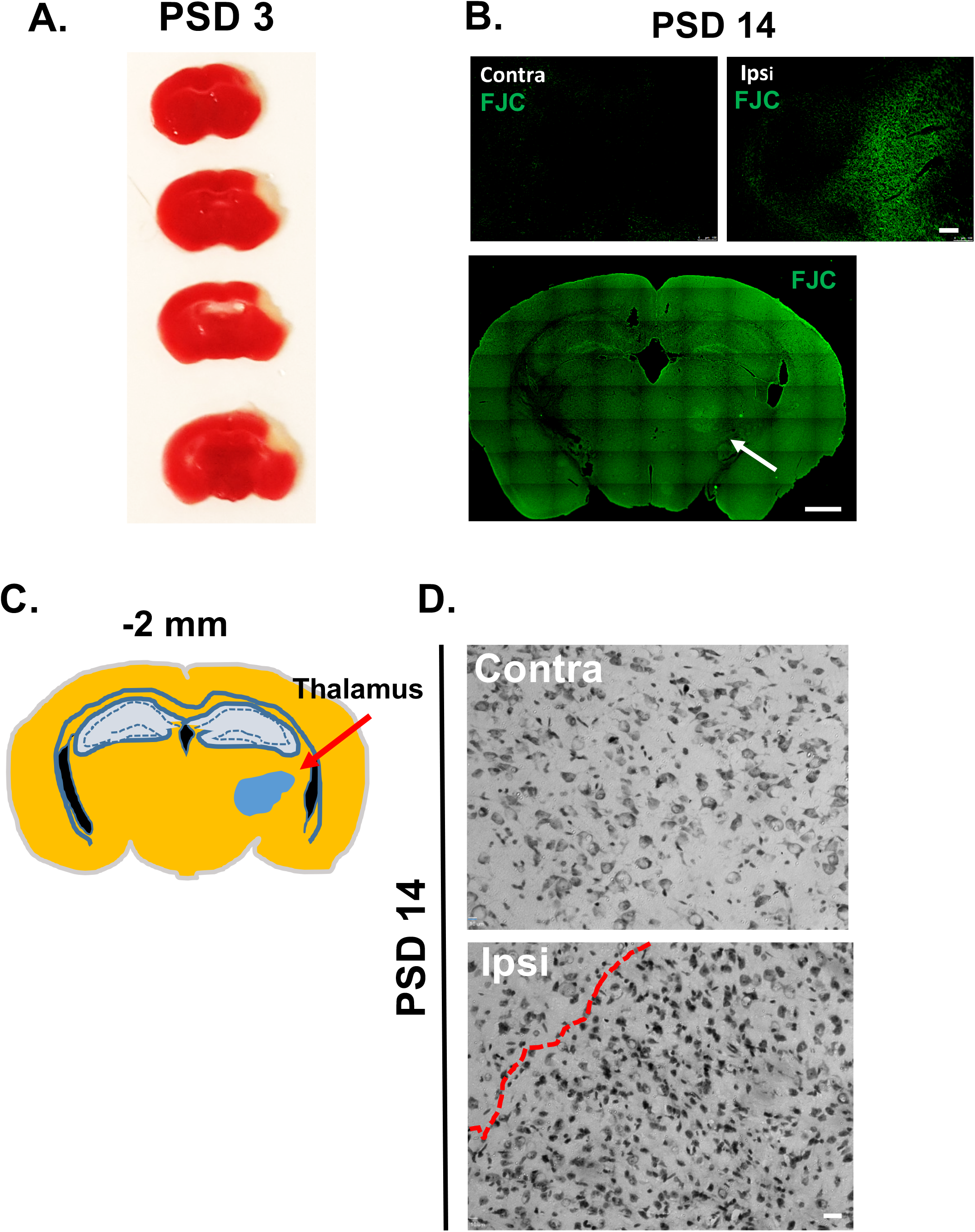
Permanent distal MCAO induces primary cortical injury with secondary injury in the thalamus. **A)** Representative TTC stained brain sections at PSD 3 showing primary injury restricted to the cortex. Distal MCAO was performed in young C57BL/6 mice. **B)** Fluoro-Jade C (FJC) staining showing degeneration of neurons in thalamus at PSD 14 (n = 5, scale bar = 100 μm in upper images, 1 mm in lower image). C) Illustration depicting the location of thalamic injury area from a brain section taken −2 mm from bregma. **D)** Cresyl violet stained sections demonstrating degenerating neurons in ipsilateral, but not contralateral thalamus at PSD 14 (n = 5, scale bar = 20 μm). Degenerating neurons are evident by their dense pyknotic nuclei.

### Cortical infarction promotes time-dependent changes in microglial activation in the thalamus

Next, evaluated the time course of microglial activation in the thalamus after cortical infarct. Region-specific microglial activation was visualized by immunostaining with an antibody against ionized calcium binding adaptor molecule 1 (IBA-1). IBA-1 is a microglia/macrophage-specific calcium-binding protein produced from the *Aif1* gene in microglia. In the brain, activated microglia express high levels of IBA-1 protein, whereas microglia in a resting state express a lower basal level. Fig. 2A shows a coronal section from a PSD 14 brain immunolabeled for IBA-1. This coronal plane (−2 mm from bregma) contains the VPM and PoM nuclei of the thalamus and the posterior extreme of the cortical infarct ^23^. Note that microglial activation is prominent and well defined within the ipsilateral thalamus (Fig. 2A); similar activation was not found in the contralateral thalamus (data not shown). In the ipsilateral thalamus, IBA-1+ microglia exhibited typical morphological changes reflective of activated microglia (amoeboid shape and hypertrophied cell body), whereas IBA-1+ microglia in the contralateral thalamus exhibited resting morphology (Fig. 2B). Examination of IBA-1 expression in brains from sham mice and PSD 3, 7, and 14 mice revealed that activated microglia were first evident in the ipsilateral thalamus at PSD 3 and became progressively more evident at PSD 7 and PSD 14 (Fig. 2C). These data show the delayed development of microglial activation within the ipsilateral thalamus, peaking well beyond the time of primary cortical infarct.

**Figure 2.**
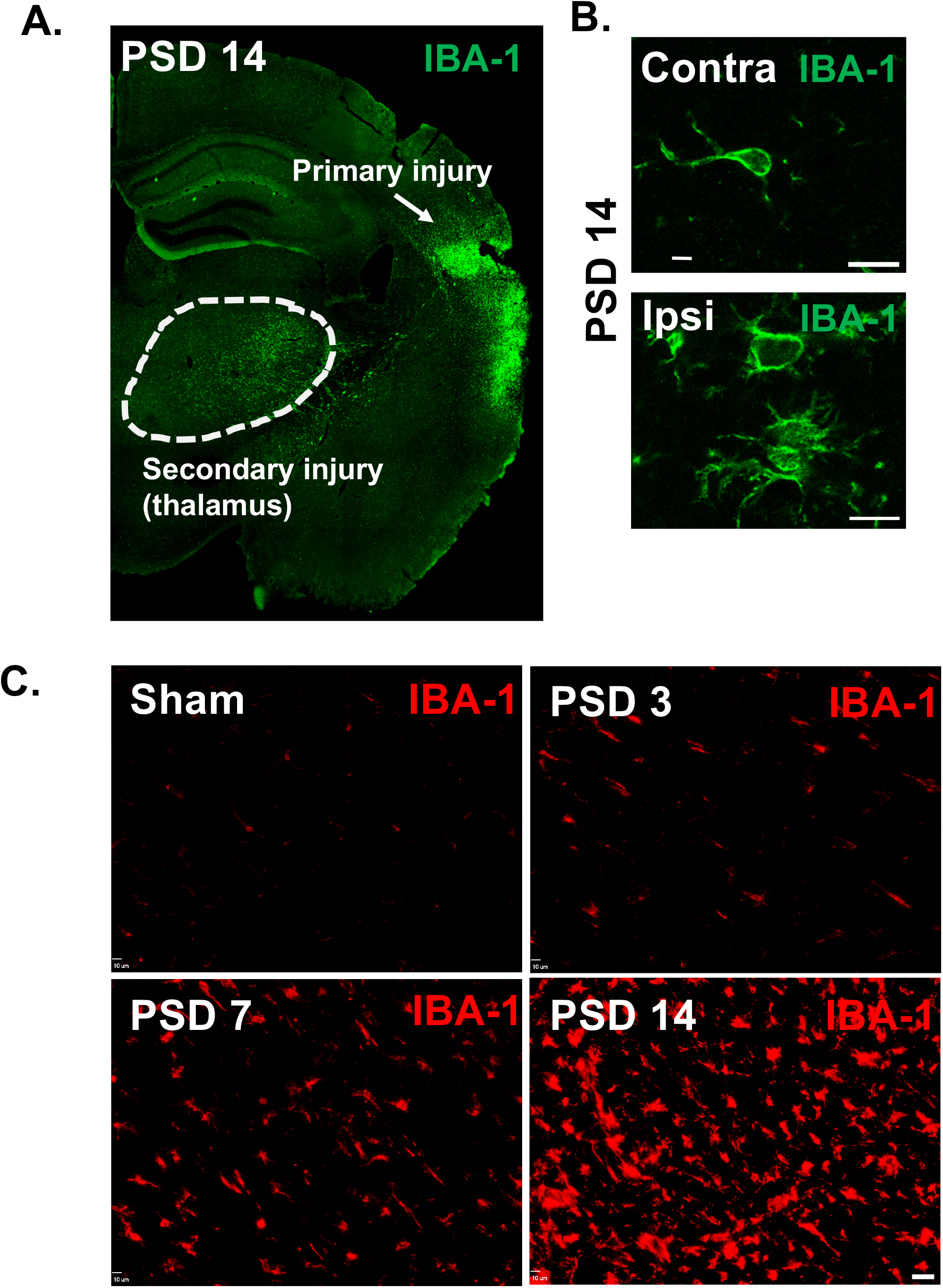
Cortical stroke induces delayed microglial activation in the thalamus. Cortical infarction was induced in young C57BL/6J mice. At 3 days, 1 week and 2 weeks after stroke, brains were isolated and sectioned. Immunostaining with anti-IBA-1 antibody was performed. Brain sections at −2 mm from bregma were used for the visualization of thalamic microglial activation. **A)** Coronal brain section showing activated MG/macrophages (IBA-1) in the posterior region of the primary injury and in the ipsilateral thalamus at PSD 14. **B)** Enlarged images showing morphology of MG/macrophages in the ipsilateral and contralateral thalamus at PSD 14 (scale bar = 10 μm). **C)** Time course of IBA-1 staining showing delayed MG/macrophage activation in the ipsilateral thalamus (scale bar = 20 μm).

### Quantitative FACS analysis demonstrates microglia in thalamus increase in number after stroke

Microglia are highly responsive to conditions found in the post-stroke brain (ischemic stress, pro-inflammatory cytokines, etc.). In the acute stroke phase (within 3 days after stroke) microglial cell number is significant reduced in the core and peri-infarct region due to massive microglial death by intensive ischemic damage and harsh hypoxic conditions ^26, 27^. In the sub-acute to chronic stroke phase (e.g. beyond PSD 3), emerging lines of evidence suggest that microglia can proliferate in damaged regions of the cortex and migrate to ischemic core, where they produce a variety of cytokines and engaged in cell debris clean up ^28, 29^. However, it remains to be elucidated whether the increased IBA-1+ cells in the thalamus that we and others ^30–32^ have shown reflects an increase in the proportion of activated microglia versus an increase in the total number of microglia. We therefore utilized flow cytometry to measure the number microglial cells from the thalamus and cortex of sham and PSD 14 brains. Cells were labeled with CD45 antibody and CD11b antibody to measure population changes in microglia as an indicator for microglial proliferation. The percentage of the microglia (CD45^int^/CD11b^+^) among the total cell population in the thalamus was increased in stroke versus sham controls (15.1% vs 6.85%, n=6) (Fig. 3A). We also found that the number of microglia counts in ipsilateral thalamus was increased compared to sham (p < 0.05, n=6). In the ipsilateral cortex, the number in microglial cells (CD45^int^/CD11b^+^) was also increased, consistent with earlier studies (Fig. 3B). These data support the possible role of either microglial migration into the region or microglial proliferation within the ipsilateral thalamus on the secondary injury after stroke.

**Figure 3.**
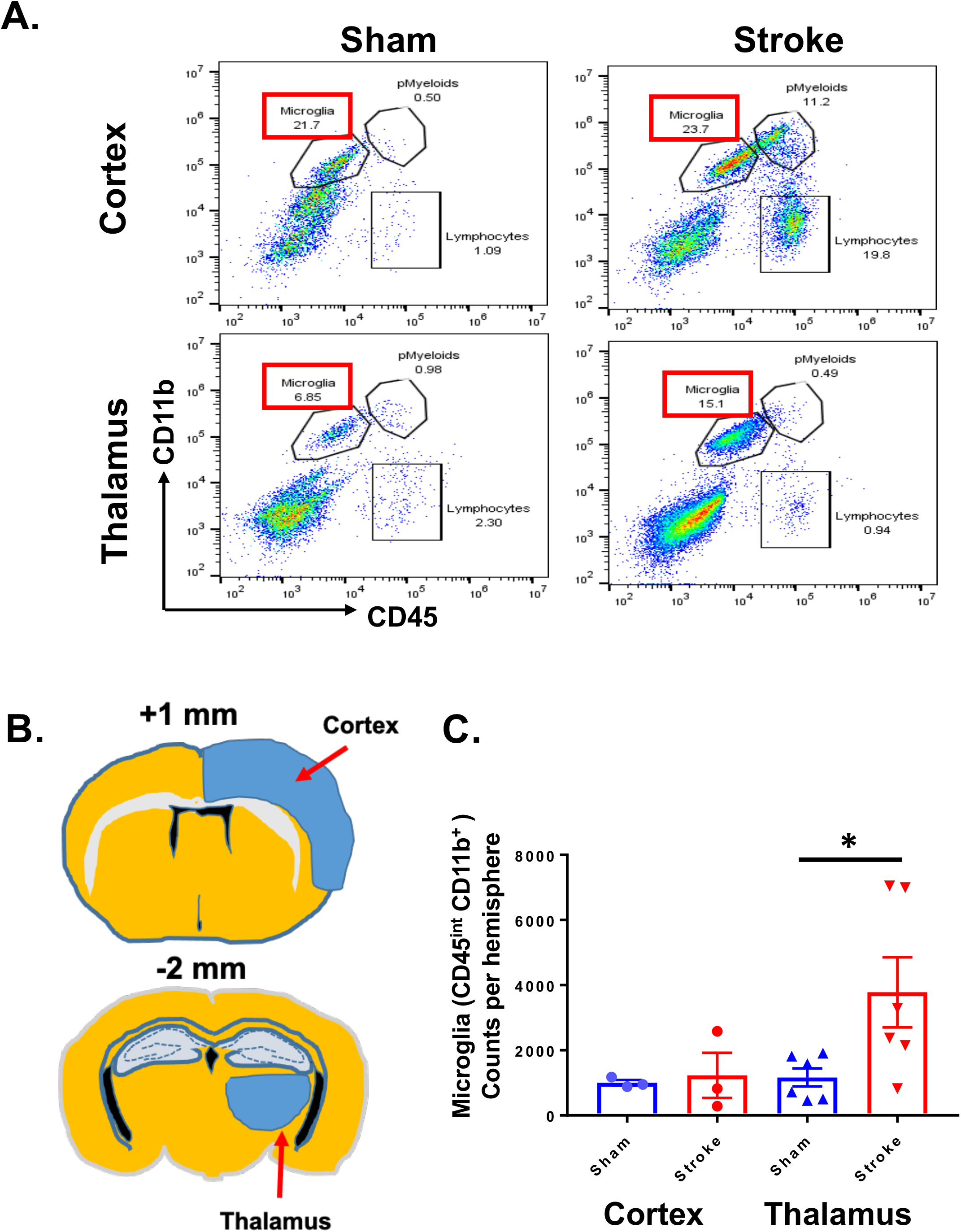
The resident microglial population increases in proportion and total number in the ipsilateral thalamus at PSD 14. At PSD 14, brains were harvested for cell cytometry. Single cell suspensions were obtained from cortex and thalamus. Cells were stained with CD45 antibody and CD11b antibody to measure the number and percentage of microglia in thalamus. **A)** Representative cell cytometry plots generated from ipsilateral cortex and thalamus from sham and PSD 14 brains. The percentage of microglia, peripheral myeloid cells (pMyeloids), and lymphocytes is indicated numerically on the plots. **B)** Illustration showing location of cortex and thalamus harvest sites from brain sections corresponding to +1 or −2 mm from bregma. **C)** Number of CD45^Int^/CD11b^+^ cells in cortex and thalamus for sham and PSD 14 brains. *P < 0.05, unpaired *t* test.

### Cortical stroke promotes recruitment of monocytes into the peri-infarct cortex as well as ipsilateral thalamus

It is known that peripheral monocytes can infiltrate the primary cortical injury and transform into IBA-1 expressing macrophages after stroke ^33, 34^. However, it is not clear if the increase in IBA-1 expressing cells in the injured thalamus reflects a similar invasion of peripheral monocytes. To determine if peripheral monocytes invasion contributes to the secondary injury mechanism, we used reporter mice for peripheral monocytes. These mice express red fluorescent protein (RFP) under the control of the C-C motif chemokine receptor 2 (CCR2) promotor (CCR2^RFP^ mice) ^35^. We performed pdMCAO in CCR2^RFP^ mice and quantified the invasion of peripheral monocytes in the cortex and thalamus at PSD 14. As expected, we found the infiltration of RFP+ monocytes within the peri-infarct cortex (Fig. 4A). Interestingly, RFP+ monocytes were also significantly increased in the ipsilateral thalamus, compared to the contralateral thalamus in 2-week post stroke brains (Fig. 4B) or thalamus of naïve mouse brain (Fig. 4C). The number of infiltrated RFP+ cells were significantly increased in ipsilateral thalamus compared to contralateral thalamus (35.3 vs 9.2 cells per mm^2^, n=3, p<0.05) after stroke (Fig. 4D). These data demonstrate that invasion of peripheral monocytes also contributes to the pathology of secondary thalamic injury.

**Figure 4.**
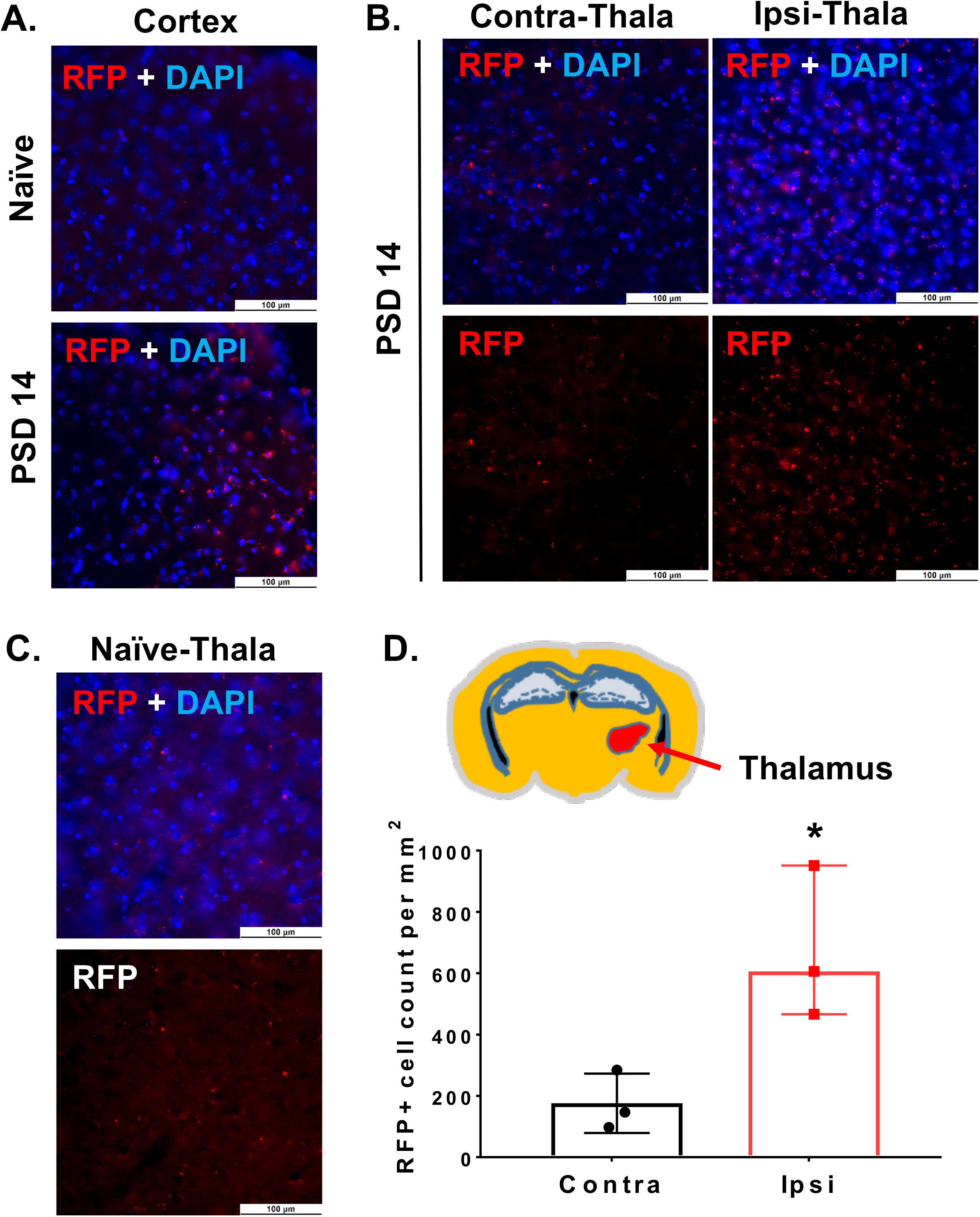
The ipsilateral thalamus demonstrates invasion of peripheral monocytes at PSD 14. CCR2-RFP reporter mice were used to track peripheral monocyte/macrophage invasion into the primary and secondary injury regions. CCR2-RFP mice underwent dMCAO and were evaluated at PSD 14. Representative images showing increased CCR2-RFP+ cells within **A)** primary injury region (cortex) and **B)** ipsilateral thalamus. The frequency of CCR2-RFP+ cells in contralateral thalamus **(B)** was similar to that of naïve CCR2-RFP reporter mice **(C)**. **D)** Quantification of RFP+ cells demonstrating increased CCR2-RFP cells in ipsilateral thalamus (versus contralateral) at PSD 14. * P<0.05, paired *t* test. Data expressed as median with 95% CI.

### Astrogliosis is evident in the thalamus after stroke

Astrogliosis is a well-documented phenomenon in primary ischemic stroke and has also been shown in secondary thalamic injury ^23, 36^. Here we evaluated the time course of astrogliosis in secondary thalamic injury. Brains were harvested at PSD 3, 7, and 14 for immunostaining to glial fibrillary acidic protein (GFAP). GFAP is an intermediate filament protein which is upregulated in activated astrocytes and used to indicate astrogliosis. At PSD 14, the VPM and PoM nuclei of the ipsilateral thalamus demonstrated pervasive astrogliosis (Fig. 5A). In contralateral thalamus, astrogliosis was not found (data not shown). The cortex, containing the posterior region of the primary infarct, also demonstrated significant astrogliosis and a dense glial scar. Earlier time points demonstrated astrogliosis in the ipsilateral thalamus by PSD 7, which continued to develop through PSD 14 (Fig. 5B).

**Figure 5.**
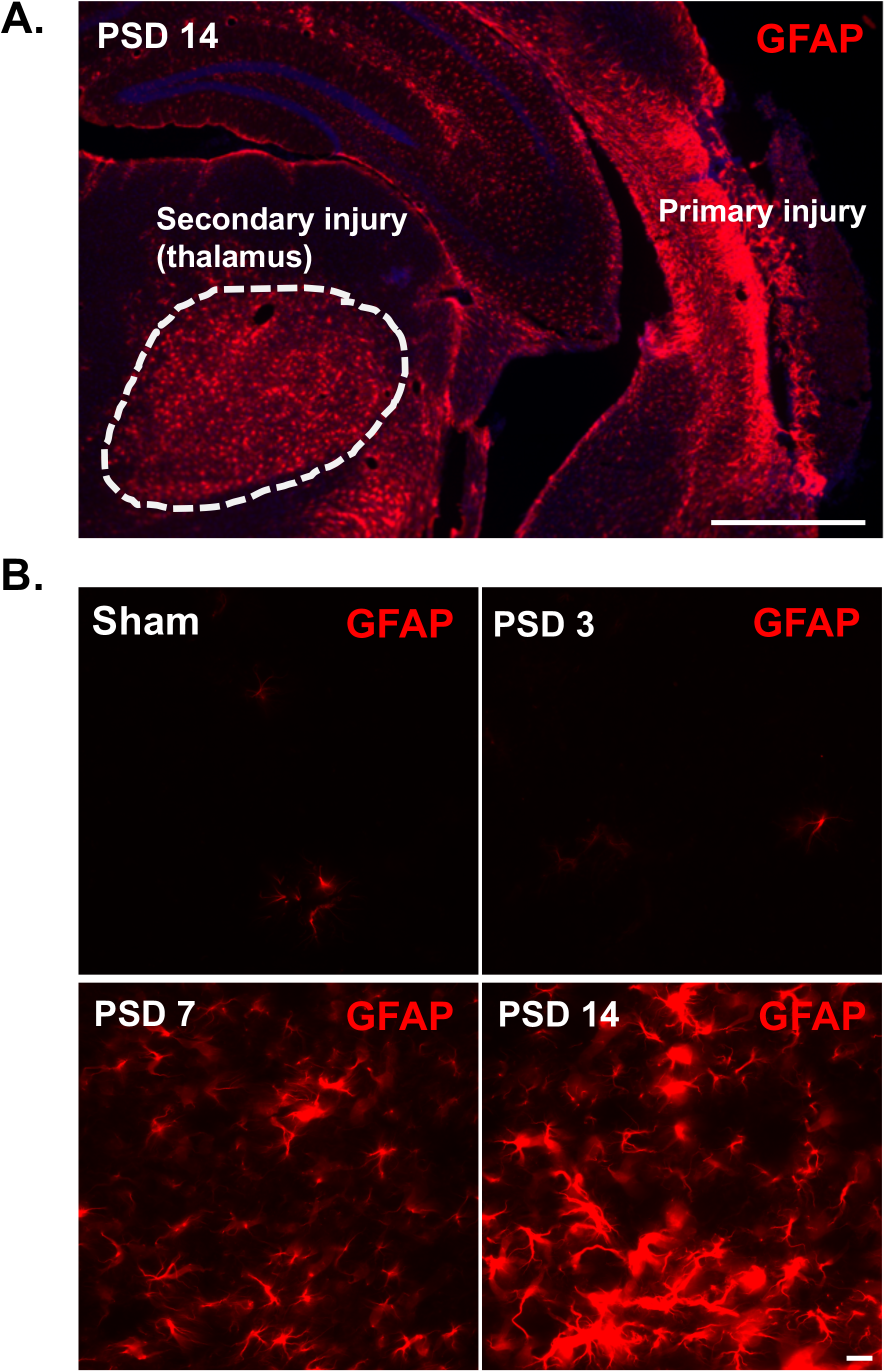
Cortical stroke induces delayed astrogliosis in the ipsilateral thalamus. Cortical infarction was induced in young C57BL/6J mice. At 3 days, 1 week and 2 weeks after stroke, brains were isolated and sectioned. Immunostaining with anti-GFAP antibody was performed. Brain sections at −2 mm from bregma were used for the visualization of thalamic microglial activation. **A)** Coronal brain section showing activated astrocytes (GFAP) in the posterior region of the primary injury and in the ipsilateral thalamus at PSD 14. A glial scar is evident in the cortical injury, but not within the thalamus (scale bar = 1 mm). **B)** Time course of GFAP staining showing delayed astrocyte activation in the ipsilateral thalamus (scale bar = 20 μm).

### Aged brains show reduced microglial activation in secondary thalamic injury

Multiple studies have shown that glial cell immune responses differ by brain region and with aging ^37, 38^. Microglia in aged brains have been shown to be more proliferative and more easily activated to pro-inflammatory cytokine and immune conditions, especially in the damaged cortex ^39, 40^. However, the effect of aging on microglial activation in secondary thalamic injury following stroke is not known. We compared pdMCAO brains at PSD14 from young and aged male mice. As expected, the young brains showed robust and pervasive microglial activation (IBA-1) across the ipsilateral thalamus (Fig. 6A). In contrast, although the aged brains showed strong IBA-1 expression in the primary cortical infarct region, the ipsilateral thalamus, showed less intense and less uniform microglial activation. Comparison of IBA-1+ area showed a significant reduction in the ipsilateral thalamus of aged mice (Fig. 6B, n= 4 or 5, p<0.05). These findings indicate that aged brains demonstrate dampened microglial activation in the thalamus, suggesting that the mechanism of secondary injury may be attenuated in aging.

**Figure 6.**
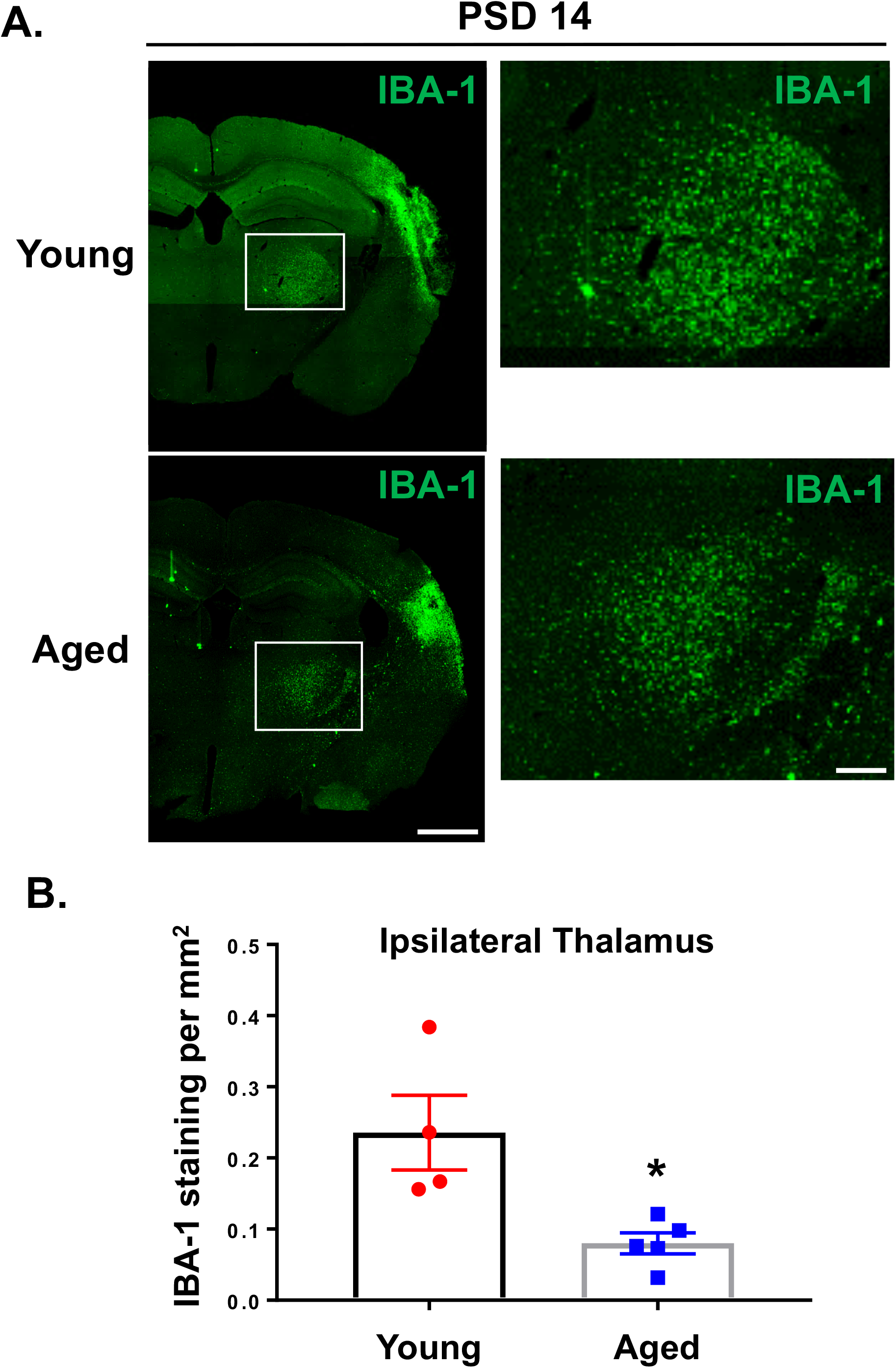
Aged mice demonstrate reduced IBA-1 expression in ipsilateral thalamus at PSD 14 compared with young mice. **A)** Representative images comparing IBA-1expression in brain of young (11-13 weeks) and aged (19-21 months) mice at PSD 14. Scale bar = 1 mm (wide image) or 250 μm (enlarged image). **B)** Quantification of total IBA-1 stained area in ipsilateral thalamus of young and aged mice. * P<0.05, unpaired *t* test.

### Aging reduces astrocytic activation in the thalamus following stroke

In order to explore how aging influences astrogliosis in the thalamus following stroke, we quantified activated astrocytes in the thalamus by GFAP immunostaining. We compared the degree of astrogliosis in the thalamus of aged and young brains following ischemic cortical stroke at PSD 7 and PSD 14. At PSD 7, astrogliosis in the thalamus of aged mice brains was significantly attenuated compared to that of young brains (Fig. 7A and 7B; n= 4/5, p<0.05). Astrogliosis at PSD14 appeared reduced in the aged brains, but did not reach statistical significance. Taken together with the reduced microglial activation in aged brains, these data suggest that aging significantly attenuates the severity of glial cell activation in secondary thalamic injury following stroke.

**Figure 7.**
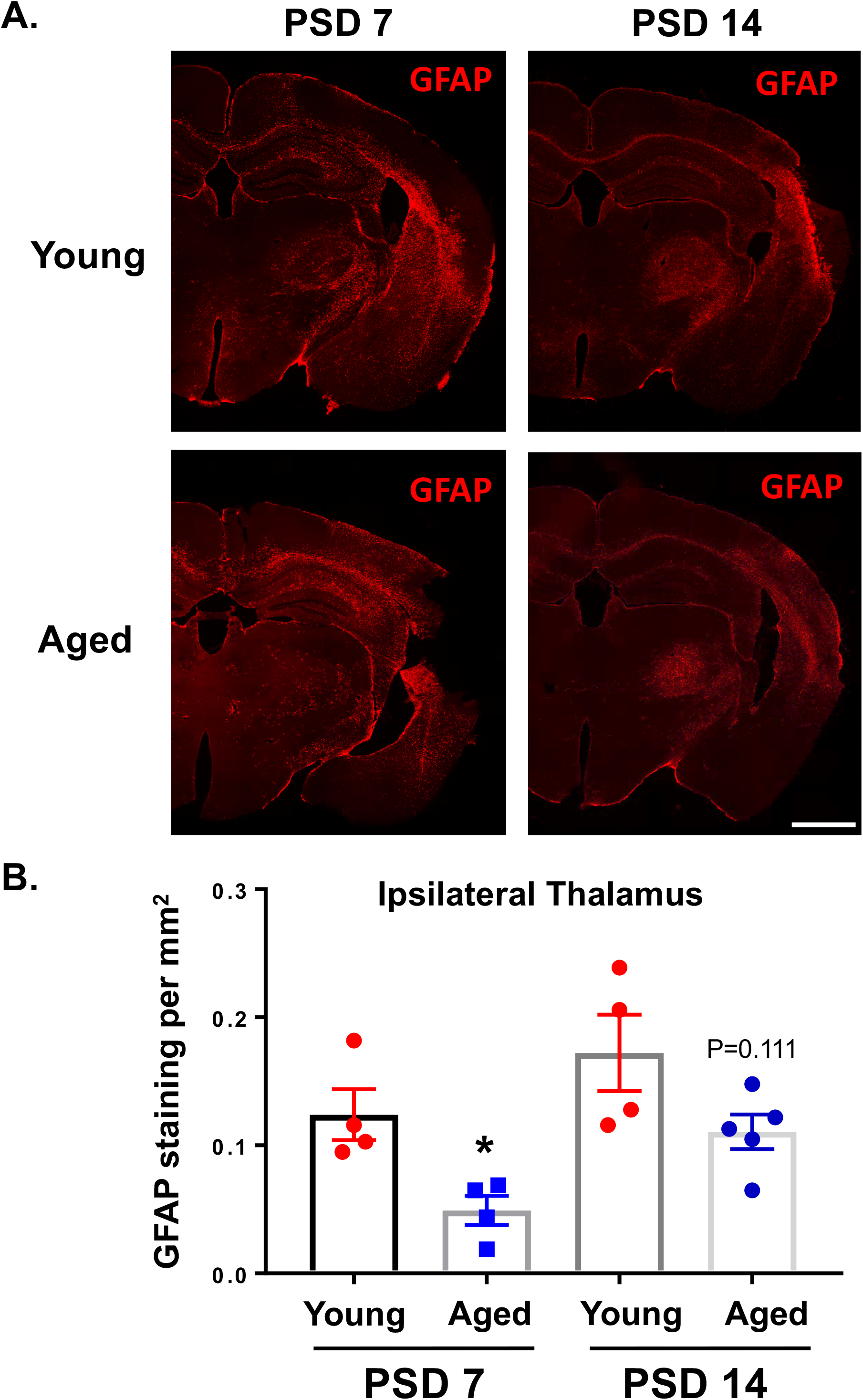
Aged mice demonstrate reduced astrogliosis in ipsilateral thalamus at PSD 14 compared with young mice. GFAP expression was evaluated at PSD 7 and PSD 14 in ipsilateral thalamus from young and aged mice. **A)** Representative images showing GFAP expression in coronal sections at −2 mm from bregma (scale bar = 1 mm). **B)** Quantification of total GFAP stained area in ipsilateral thalamus of young and aged mice. * P<0.05, unpaired *t* test. (P=0.111 for young versus aged at PSD 14).

### Glial scar like astrogliosis is observed in the thalamus of 2-year post-stroke brain

We have shown here and previously ^23^ that, unlike cortical astrogliosis (which includes diffuse gliosis *and* glial scar formation), the astrogliosis that occurs in the ipsilateral thalamus retains a more uniform pattern and lacks obvious scar characteristics. However, given the progressive development of astrogliosis that was observed over time, it was important to define the long-term characteristics of gliosis in this specific type of injury. Up to this point, glial scar formation is typically reported for primary injury to the cortex (ischemic and traumatic injury) or spinal cord injury. Therefore, to determine the long-term characteristics of astrogliosis in secondary thalamic injury, we compared GFAP immunoreactivity in mice at 6 weeks and 2 years post-stroke (young mice at the time of stroke). Astrogliosis in the ipsilateral thalamus at 6 weeks post-stroke was similar to that at PSD 14 (Fig. 8A). In particular, gliosis was widespread throughout the thalamic region, but without apparent glial scar characteristics. Note that glial scar was still evident at the site of primary infarct in the cortex, particularly in more anterior sections. At 2 years post-stroke, astrogliosis was still prominent in the ipsilateral thalamus (Fig. 8B). Interestingly, the gliosis had incorporated glial scar-like properties, including a dense scar core with surrounding reactive astrocytes (Fig. 8C). To our knowledge, this represents the first report of glial scar in secondary thalamic injury. Astrogliosis was broadly elevated in the aged brains, particularly in the fiber tracts (e.g. corpus callosum, fimbria, internal capsule), hippocampus, and caudoputamen. However, note that gliosis in the contralateral thalamus was negligible compared to the ipsilateral thalamus. These data show that the astrogliosis that occurs as a result of secondary thalamic injury continues to evolve for months or possibly years, with the potential to develop scar-like features. In addition, the astrogliosis that is produced in secondary thalamic injury is extremely long-lasting, and possibly permanent.

**Figure 8.**
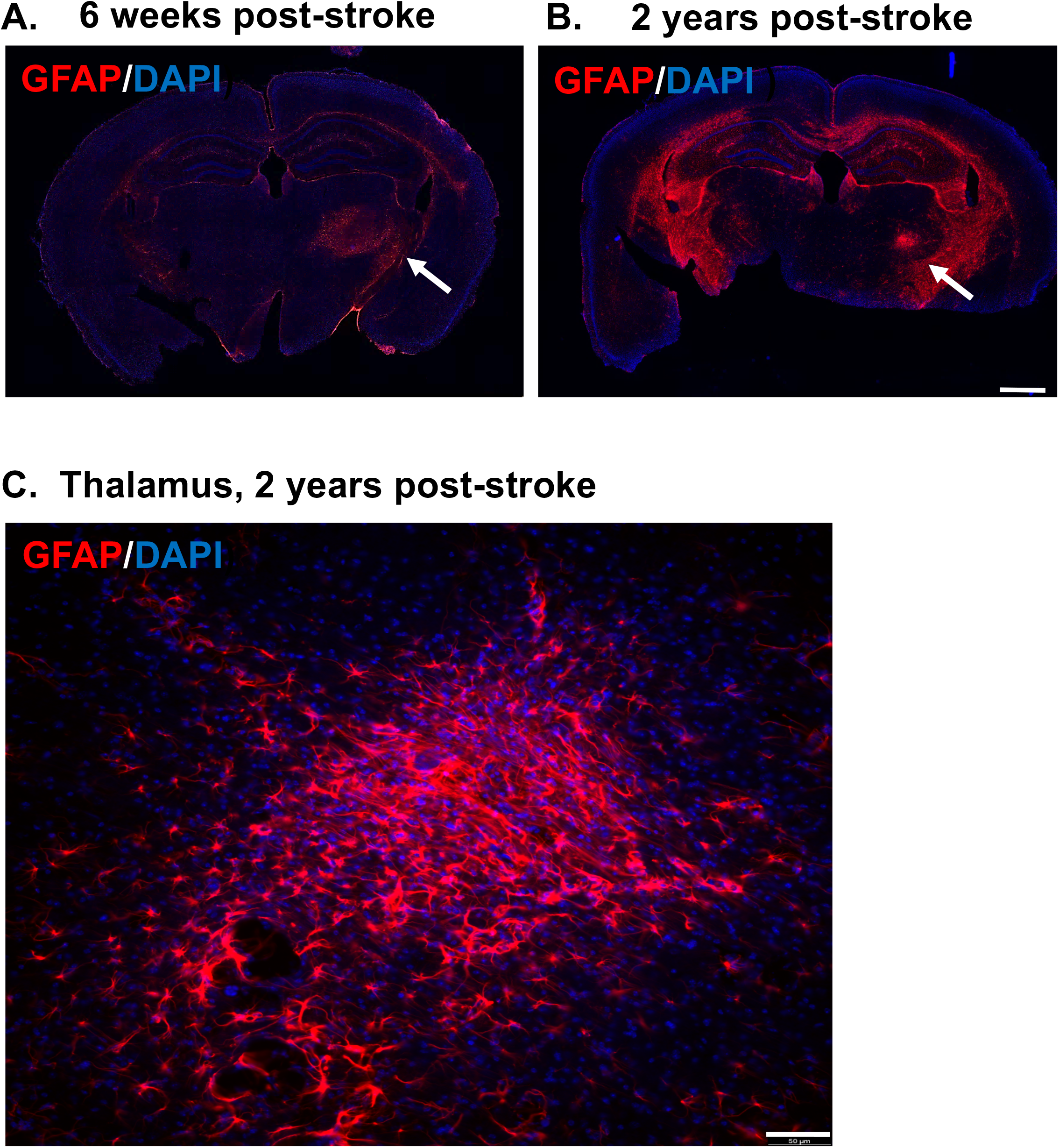
Astrogliosis in ipsilateral thalamus is long-lasting and can eventually develop glial scar-like characteristics. Young mice were subjected to dMCAO and then allowed to recover for 6 weeks or two years. At 6 weeks or two years post-stroke, brains were evaluated for astrogliosis (GFAP). Brain sections showing astrogliosis at 6 weeks **(A)** or two years following stroke **(B)**. Scale bar = 1 mm. **C)** Enlarged region of thalamus from two year post-stroke brain showing glial scar-like characteristics. Scale bar = 50 μm.

### Thalamic gliosis can be modifiable with NMDA receptor antagonist following stroke

Secondary thalamic injury is thought to be mediated by the mechanisms of excitotoxicity, in part through NMDA pathways ^41^. Therefore, we lastly sought to determine if secondary thalamic injury or gliosis could be reduced by treatment with a NMDA receptor antagonist (memantine). Memantine is an FDA-approved drug that has been reported to ameliorate primary cortical damage in preclinical study after stroke ^42–44^ and has been proven safe in humans. In the clinical setting, memantine has been widely used for treating Alzheimer’s disease (AD) patients as it has been proven to reduce NMDA mediated neurotoxicity and inflammation, thus reducing the symptoms of AD. Many studies have already shown that memantine reduces primary infarction and improves functional outcome in stroke animal models. To determine if memantine treatment can reduce secondary thalamic injury, we administered memantine (or vehicle) following pdMCAO in young mice. Memantine was delivered at 4 hours post-stroke (100 mg/kg, ip) and again at 24 hours post-stroke (50 mg/kg, ip). Brains were isolated at either 3 days (for TTC measurement of cortical infarct volume) or 14 days (for quantification of thalamic gliosis). At PSD 3, the memantine-treated mice showed a non-significant reduction of infarct volume compared with vehicle treatment (MEM;18.4 ± 5.32 vs. VEH; 24.86 ± 1.52 mm^3^, n=6/5, *p*=0.3131) (Fig. 9A, B). However, at PSD 14, the area of GFAP+ staining was significantly reduced by memantine treatment (Fig. 9C, P<0.05, n=9). In addition, the area of IBA-1+ staining approached a significant reduction with memantine treatment (Fig. 9D, P<0.05, n=9). Taken together, these data indicate that secondary thalamic injury can be modified by pharmacological approaches targeting NMDA receptor activation (e.g. excitotoxicity) in the post-stroke period.

**Figure 9.**
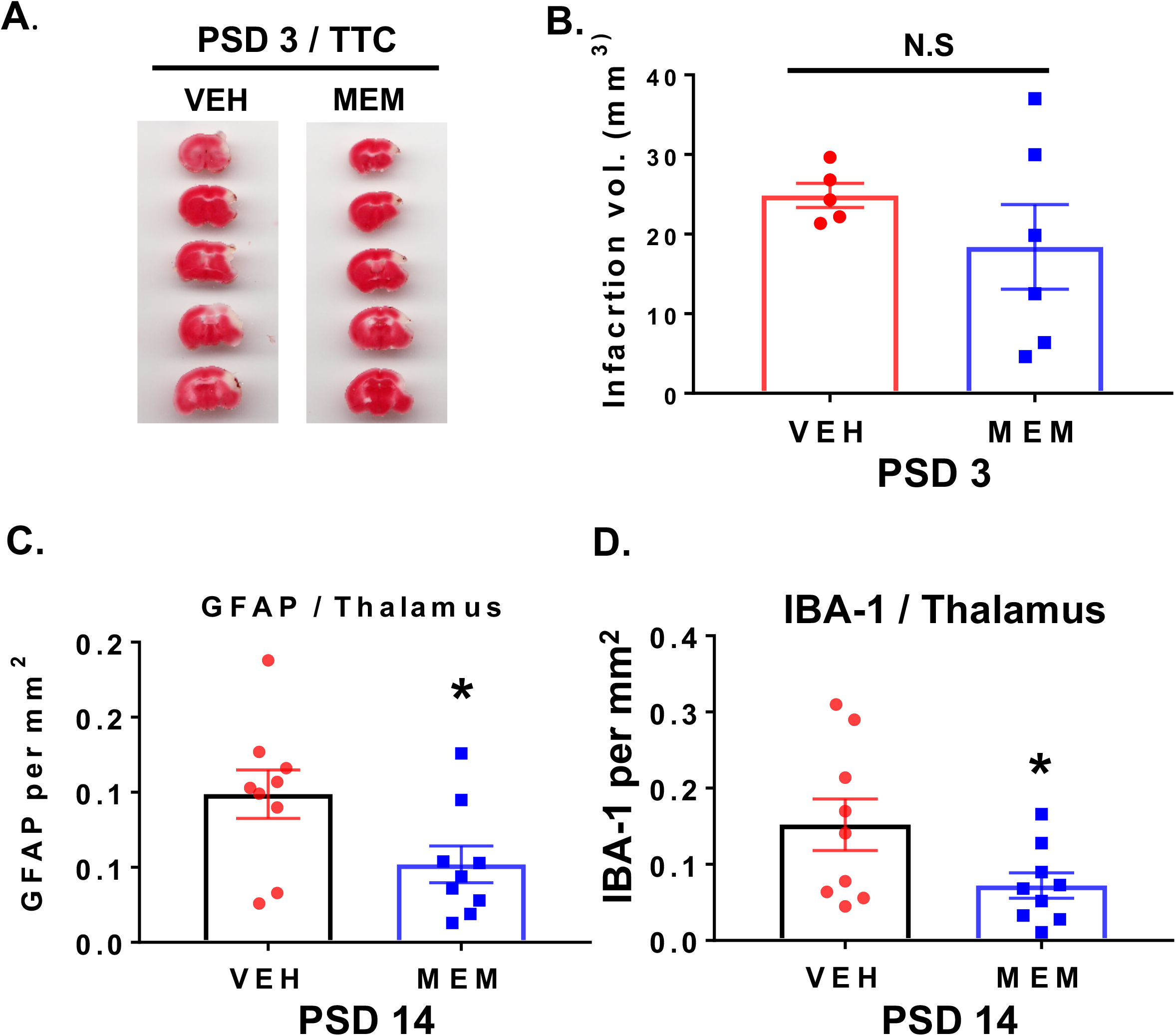
Memantine, an NMDA antagonist reduced glial activation in thalamus after stroke. Post-stroke treatment with memantine, an NMDA receptor antagonist, reduces gliosis in ipsilateral thalamus. Representative images showing TTC staining **(A)** and infarct volume quantification **(B)** at PSD 3 for memantine and vehicle treated mice. Primary cortical infarct volume was not significantly changed with memantine treatment. **C)** Thalamic area demonstrating GFAP expression was significantly reduced at PSD 14 with memantine treatment. * P < 0.05, unpaired *t* test. **D)** Thalamic area demonstrating IBA-1 expression approached a statistical reduction with memantine treatment. * P < 0.05, unpaired *t* test.

## DISCUSSION

In this study, we used a mouse model to explore the temporal and cellular changes associated with secondary injury in the thalamus following stroke in distant brain regions. Using this model system, we made the following novel discoveries. First, we showed that the gliosis response in the thalamus consists of a significant expansion of the microglia/macrophage cell population, which is due in part to the invasion of peripheral monocytes. Second, we showed that the thalamic gliosis response is attenuated in aged mice. Third, we demonstrated that astrogliosis in the ipsilateral thalamus is long-lasting (possibly permanent) and eventually develops glial scar-like characteristics. Fourth, post-stroke treatment with NMDA receptor blocker, memantine, attenuates gliosis in the thalamus. We expand on these new findings below.

### The gliosis response in the thalamus consists of a significant expansion of the microglia/macrophage cell population, which is due in part to the invasion of peripheral monocytes

It is well-established that CNS injury initiates the intrinsic programs for microglia to act as a first response immune cell in the brain. In this process, activated microglia play a critical role in sensing neuro-inflammation and releasing chemokines and cytokines (pro- and anti-inflammatory), which spread from the damaged area to the spared but active brain area ^45, 46^. A large body of evidence on how microglial phenotypes are altered by ischemic injury explains the regional diversity and modes of microglial polarization from M0 to M1 or M2 in neurodegenerative disease models, in response to the severity and types of injuries ^47, 48^. During the acute stroke phase, the ischemic stress causes microglia in the core and penumbral regions to undergo apoptosis and necrosis, thus resulting in the decrease in microglial cell number ^26, 27^. Subsequently, the microglial cell population can return through proliferation of remaining MG or migration of MG from nearby regions ^49–51^. In addition, CCR2^+^ peripheral monocytes can infiltrate the injury region and then differentiate into macrophages or microglia-like cells ^34, 52, 53^. The differentiation of the macrophages is affected by the local environment as well as the type and time course of injury ^54^. Whether peripheral monocytes invade or contribute to a microglia/macrophage population in secondary injury is not currently known.

We used a combination of immunostaining, flow cytometry, and CCR2 reporter mice to examine the temporal and cellular origin of gliosis in the thalamus after stroke. Similar to what others have shown, the cortical infarct induced delayed and progressive microgliosis (IBA-1^+^ cells) within the VPN/PoM nuclei of the ipsilateral thalamus ^30, 55^. Using flow cytometry, we found that the proportion and absolute number of CD45^int^/CD11b^+^ MG was significantly increased in PSD 14 thalamus. The CD45^int^/CD11b^+^ cells are reflective of the total MG population, not simply the activated MG (i.e. IBA-1^+^ cells). The four-fold increase in total MG in the ipsilateral thalamus (versus sham) indicates a substantial expansion of this population above pre-injury levels. In comparison, the absolute number and proportion of CD45^int^/CD11b^+^ MG in cortex at PSD 14 was similar to sham controls. The cause of this difference between cortex and thalamus was not determined, but could reflect a less extensive (or no) initial MG depletion following thalamic injury or a different time course of MG depletion/repopulation. In addition, the proportional value of MG in the cortex sample reflects a dilution effect from the increase in peripheral myeloid cells and lymphocytes. Therefore, to more specifically evaluate the region and injury-specific potential for peripheral monocyte invasion, we performed pdMCAO in CCR2 reporter mice. These mice express RFP in CCR2-expressing monocytes and monocyte lineage cells (e.g. monocyte-derived macrophages), but not in resident CCR2-negative macrophages. As expected, the cortex demonstrated the presence of RFP^+^ monocyte/macrophages at PSD 14. Interestingly, the ipsilateral thalamus also demonstrated a significant increase in RFP^+^ cells, indicating that peripheral monocytes do invade the thalamus following secondary injury. To our knowledge, this represents the first report of peripheral monocyte infiltration as part of the secondary injury mechanism. In primary stroke injury, circulating CCR2^+^ monocytes have been shown to be recruited to ischemic regions from the leptomeninges surrounding the brain parenchyma and then along penetrating arterioles ^34^. However, given the greater distance of the thalamus from the leptomeninges, monocyte influx from the blood circulation may be more likely in this brain region. Nevertheless, regardless of how the monocytes ultimately arrive, these findings suggest that peripheral monocytes can contribute to the expansion of the IBA-1^+^ MG/Macrophage population in secondary injury in the thalamus. Future studies are warranted to determine the signaling mechanisms and route of entry for recruited peripheral monocytes, the time course of monocyte infiltration, and the role of this monocyte-derived population in the mechanism of secondary injury or repair.

### The thalamic gliosis response is attenuated in aged mice

The risk of ischemic stroke is greatly increased in aging, showing a doubling of stroke incidence for each decade after 55 years of age ^56^. Therefore, to increase the translational value of these studies, we performed a subset of studies in aged mice (18-22 months). This age range in mice corresponds roughly with that of humans in the 6^th^ to 7^th^ decade of life ^57, 58^.

Aging is associated with a number of pathological changes in MG and astrocytes, two glial cell types which play important roles in both brain support and response to injury ^59, 60^. Much of the age-dependent increase in neuroinflammation has been linked to changes in microglia, which exhibit both morphological and molecular changes consistent with an enhanced inflammatory profile ^60^. These “primed” phenotype MG in the aged brain demonstrate increased expression of antigen presentation and antigen receptor genes and demonstrate an exaggerated inflammatory response to immune stimuli ^61^. Astrocytes also demonstrate age-dependent impaired support function and exaggerated responsiveness to injury. Aged astrocytes produce lower amounts of trophic factors, such as VEGF and Wnt3, exhibit reduced glutamate uptake and clearance, and contribute to accelerated glial scar formation ^59^. In general, aging leads to a brain environment in which glial support function is reduced and responsiveness to inflammatory stimuli is amplified. However, evidence for how aging affects the MG and astrocytes response in secondary injury is currently lacking.

We showed that aged mice, despite the formation of a classic glial scar in the cortex, exhibited reduced astrogliosis and microgliosis in the thalamus following stroke. Given the enhanced inflammatory responsiveness in the aged brain, this outcome was contrary to our initial prediction. However, a possible explanation for the reduced gliosis response may involve age-dependent cell senescence, as some reports have shown MG senescence and hypoactivation in aging ^60, 62, 63^. In addition, MG in the aged CNS have also been reported to have reduced potential for replication due to telomere shortening and demonstrate reduced proliferation following injury ^63, 64^. The combination of reduced activation and reduced proliferation, could account for the reduced IBA-1+ population in the ipsilateral thalamus and a reduced source of inflammatory stimulation of astrocytes. To our knowledge, our study represents the first report of how aging specifically affects the pathology of secondary thalamic injury.

### Astrogliosis in the ipsilateral thalamus is long-lasting (possibly permanent) and eventually develops glial scar-like characteristics

The astrogliosis response in the thalamus consisted of a diffuse distribution of GFAP^+^ astrocytes throughout the VPM and PoM nuclei, becoming first evident between PSD 3 and 7. The astrogliosis in thalamus differed from that in the primary cortical injury region in that a glial scar did not form around the injury within the first two weeks, or even 6 weeks. However, at two years after injury, a dense highly-GFAP^+^ cell cluster developed within the ipsilateral thalamus. This cluster demonstrated the dense mesh-like arrangement of activated astrocytes characteristic of a glial scar ^65^. The contralateral thalamus was notably lacking in gliosis, thus supporting the causative role of the primary stroke injury on the enduring gliosis in the ipsilateral thalamus. While the aged brains did show increased astrogliosis in a variety of brain regions (particularly in the fiber tracts), it showed bilateral distribution.

The formation of a glial scar is a common feature in primary stroke injury, as well as several other types of severe injury in the CNS ^65–68^. The glial scar is composed mainly of hypertrophic astrocytes which characteristically express increased levels of GFAP ^65^. The scar typically forms at the boundary between necrotic and healthy tissues and appears to support tissue repair mechanisms in the CNS ^69, 70^. However, how the scar either impedes or supports axon regrowth is still hotly contested ^69^. The cause or purpose of glial scar formation in the thalamus perplexing. The scar in this region appears to take many months (or possibly years) to develop; thus, the purpose appears to be inconsistent with that of a protective barrier to support repair. Given that the scar was shown in mice two years following stroke, it is possible that the slow development of the scar might be related to the changing environment within the aging brain. In light of the lengthy time over which this secondary injury develops, it will be important to evaluate the potential to either reduce (i.e. therapeutic) or worsen (i.e. environmental stress, etc.) secondary injury at significantly later time points.

As a result of the advent of new technologies, such as comprehensive single cell RNA sequencing and large-scale transcriptome analyses, we are gaining a better appreciation of the clear heterogeneity in astrocytes in CNS. This heterogeneity, which is driven by physiological and pathological conditions as well as aging, accounts for specialized functions of these cells in different brain regions, circuits and networks ^71, 72^. Thus, each type of astrocyte can play a unique role in sensing and responding to the diverse cellular signaling, which occurs in different steps of inflammatory responses. The formation of the glial scar is an example of specialized astrocytic function in the context of stroke. It is well established that the glial scar, which forms around the infarcted area, contributes to the inhibition of the spread of inflammation to nearby healthy tissue, the production of neurotrophic factors in repair/remodeling, and the generation of a scaffold for facilitating axonal regrowth ^73, 74^. However, as noted above, the injury within the ipsilateral thalamus showed broad astrocyte activation, but no evidence of glial scar formation – even through six weeks after stroke. The reason for the different gliosis pattern may reflect different types of injury (ischemic vs. excitotoxic) as well as different subtypes of glial cells in the respective regions. A recent study using scRNAseq revealed seven distinct types of astrocytes with region-specific distributions in the brain ^71^. The authors found that two major sub-types of astrocyte populations (telencephalon astrocytes) are distributed in the cortex, striatum, amygdala, and hippocampus. However, they showed that these sub-types are absent in the thalamus and midbrain, highlighting the heterogeneity of astrocytes in the cortex and thalamus.

### Post-stroke treatment with NMDA receptor blocker, memantine, attenuates gliosis in the thalamus

The thalamus plays many critical roles in the brain, and serves as a vital relay station between different brain regions (e.g. cortex and brainstem). However, as a result of the high connectivity of this brain center, the thalamus is also vulnerable to secondary injury from multiple brain regions. Remote secondary thalamic injury has been observed in rodent stroke models as well as in human stroke pathology. Furthermore, thalamic dysfunction is associated with worsened outcome and recovery following stroke ^7, 9, 75^. It is significant to note that the development and progression of this secondary injury appears to be modifiable in experimental models. We have previously shown that thalamic injury (neuronal loss and gliosis) can be reduced with post-stroke hypothermia ^23^, while others have shown similar benefit with low oxygen post conditioning ^41^. In the other direction, others have shown that the neuronal loss can be enhanced by 4 weeks of post-injury chronic stress ^24^. From a clinical perspective, the slow development and modifiable nature of secondary injury is encouraging, as it means this type of injury is amenable to delayed therapeutic interventions.

Memantine is an NMDA receptor antagonist that has been previously demonstrated to reduce primary infarction in experimental models of both permanent and transient stroke models ^42–44^. This drug is also currently used clinically in the treatment of Alzheimer’s disease symptoms ^76^. Although the mechanism of secondary injury is not fully elucidated, it does appear that glutamate-mediated excitotoxicity plays a key causative role ^77^. Whether post-stroke treatment with memantine could reduce the secondary thalamic injury and gliosis has not previously been studied.

Our studies indicated that treatment with memantine at 4 and 24 hours after stroke results in reduced gliosis in the ipsilateral thalamus. At PSD 14, astrogliosis was significantly reduced and microgliosis approached significant reduction (P=0.056) in memantine treated mice. The primary cortical infarct (evaluated at PSD 3 in a separate cohort) was not significantly reduced with memantine treatment. This significant effect on secondary injury with lack of significant effect on primary injury suggests that memantine is acting to disrupt the progression secondary, similar to our findings with hypothermia (Cao 2017). The timing of memantine for this initial study was chosen in part on clinical practicalities. Future studies would be required to determine the effective treatment window for memantine in reducing secondary injury.

### Limitations

Our studies were performed exclusively in male mice. However, it has been shown that there are sex differences in microglial and astrocyte responses to ischemic injury, in both inflammatory response and cell survival following ischemic stress ^78–80^. Future studies will be required to determine how secondary thalamic injury is affected by sex hormones or genetic sex.

### Summary

Secondary injury can be produced in distal brain regions via interconnecting neuronal pathways with primary sites of ischemic injury. Relay centers, such as the thalamus, may therefore be particularly vulnerable to this type of injury. We found that cortical stroke induces progressive and lasting secondary injury in the ipsilateral thalamus. This thalamic injury involves initial gliosis, followed by neuronal loss in the VPN and PoM nuclei. Over a period of several months, the gliosis in the thalamus eventually develops characteristics of a glial-scar. In aged mice (at the time of stroke), gliosis in the thalamus is attenuated. We also showed for the first time that peripheral monocytes invade the injured thalamus and can therefore contribute to the expansion of the activated microglia/macrophage population in secondary injury. Lastly, we showed that post-stroke treatment with an NMDA receptor antagonist results in reduced gliosis in the thalamus. These findings provide an important foundation for future studies on the molecular mechanisms of thalamic injury and for strategies aimed at reducing secondary brain injury after stroke.

## Methods

### Animals

All procedures were performed in accordance with NIH guidelines for the care and use of laboratory animals and were approved by the Institutional Animal care and use committee of the University of Texas Health Science Center. Male C57BL/6J mice (11-14 weeks old and 18-22 month old) were used for all experiments. All animals were housed in the animal care facility and had access to water and food *ad libitum* at University of Texas Health Science Center.

### Permanent distal middle cerebral artery occlusion (pdMCAO) model

We used male C57BL/6 mice, between 11–14 weeks (young) or 18-22 months (aged). Permanent focal cerebral ischemia was induced by permanent occlusion of the right distal middle cerebral artery (MCA) using electro-coagulator (Accu-temp). The distal MCA was accessed via a craniotomy and permanently occluded just proximal to the anterior and posterior branches. Mice were anesthetized with isoflurane (4% induction and 2% maintenance in airflow) during the surgery. Body temperature (rectal probe) was maintained at 37 °C during surgery. Bupivacaine (1ml/kg of 0.25% solution, s.c.) was injected prior to skin incision for pain management. A skin incision was made between the ear and eye and the temporal muscle was detached from the skull to locate the MCA beneath the transparent skull. A small craniotomy was then generated over the MCA using a micro-drill to provide access to the artery. An Accu-temp® variable low temperature cautery was used to permanently ligate the MCA. After surgery, mice received heat support for 2 hours. Sham controls underwent the same procedure, except for the ligation of the MCA.

### Brain sample preparation and immunostaining

Mice were euthanized at 3 or 14 days after stroke. Mice were transcardially perfused with heparinized PBS, followed by 4% PFA (paraformaldehyde in PBS). Brains were harvested and then fixed for an additional 24 hours in 4% PFA at 4°C. Brains were then transferred to 30% sucrose solution in PBS for 24h hours prior to generating 30 μm coronal sections (Micron HM 450, Thermo fisher scientific, MA, USA). For immunostaining, sections were washed with PBS, incubated with blocking buffer (10% goat serum, 0.3 % Trion X-100 in PBS), and then incubated overnight at 4°C with the following primary antibodies: Rabbit anti-IBA-1 antibody (1:200) (Wako pure chemical, Japan), mouse anti-GFAP antibody-cy3 (1:500) (MilliporeSigma, MO, USA). For detection of the IBA-1 antibody, we used either donkey anti-rabbit IgG-Alexa 594 or 488 (1:200, Thermo fisher scientific, MA, USA). Sections were incubated with DAPI (4′,6-diamidino-2-phenylindole) to label nuclei. Images were obtained using a Leica TCS SPE confocal system and a Leica DMi8 fluorescence microscope system (Leica biosystem, IL, USA). Stitched images were generated with Leica LAS X software. Image analysis was performed using ImageJ software (National Institutes of Health).

### Identification of CCR2+ cell in brains

pdMCAO was performed on CCR2^RFP^ mice (Jax017586 - B6.129(Cg)- CCR2<tm2.1Ifc>/J, male) ^35^ in which red fluorescent protein is constitutively expressed under the chemokine (C-C motif) receptor 2 (CCR2) promotor. At 2 weeks after surgery, brains were isolated and sectioned as described above. Sections were incubated with DAPI and then imaged and analyzed as described above.

### Flow cytometry

Mice (sham and stroke group) were anesthetized with Avertin® (2,2,2-Tribromoethanol, Sigma, MO, USA) and transcardially perfused with phosphate-buffered saline (PBS) for 5 min. Brains were isolated and then immediately separated into cortex and thalamus. Brain tissue was then placed in complete Roswell Park Memorial Institute (RPMI) 1640 medium, followed by mechanical and enzymatic digestion with 150 μL collagenase/dispase (1 mg/mL) and 300 μL DNAse (10 mg/mL; both Roche diagnostics, IN, USA) for 45 min at 37 °C with mild agitation. The tissue was triturated by pipette, cleared of myelin and cell debris (cell debris removal kit, Miltenyi Biotec, Germany), and then filtered through a 70 μm separation filter. Cells were washed and blocked with mouse Fc Block (eBioscience, CA, USA) prior to staining with primary antibody conjugated fluorophores: CD45-eF450 (# 48-0451-82, eBioscience, CA, USA), CD11b-AF700 (# 101222, Biolegend, CA, USA). All the antibodies were purchased from Biolegend/eBioscience. For live/dead discrimination, a LIVE/DEAD Fixable Aqua Dead Cell Stain Kit was used according to manufacture instructions (Thermo Fisher Scientific, MA, USA). Cells were briefly fixed in 2% PFA. Data was acquired on a CytoFLEX cytometer (Beckman Coulter, CA, USA) and analyzed using FlowJo (BD biosciences, OR, USA). No less than 100,000 events were recorded for each sample. Cell type-matched fluorescence minus one (FMO) controls were used to determine the positivity of each antibody.

### Data Analysis

All the data were expressed as mean ± SEM, except Figure 4D (median with 95% CI). GraphPad Prism (version 7.03, San Diego, CA) was used to analyze and plot the data. Student’s unpaired t-test and paired t-test were used for two group comparisons. A p-value < 0.05 was considered statistically significant

## Acknowledgements

This project was funded by NIH NS096186 (S.P.M) and NS 094280 (S.P.M).

## Author Contributions

S.P.M. and G.S.K. conceived the experiments. G.S.K., J.M.S., A.A.M., T.W. performed the experiments. G.S.K., S.P.M., J.L. and F.L. analyzed the results. S.P.M., G.S.K., M.G.G. and J.W.M. discussed the results. S.P.M. and G.S.K. made the figures and wrote the manuscript. All authors reviewed the manuscript.

